# Phosphorylation by Aurora B kinase regulates caspase-2 activity and function

**DOI:** 10.1101/2020.03.05.978056

**Authors:** Yoon Lim, Dylan De Bellis, Jarrod J. Sandow, Luisa Capalbo, Pier Paolo D’Avino, James M. Murphy, Andrew I. Webb, Loretta Dorstyn, Sharad Kumar

**Author notes:** Senior authors (Loretta Dorstyn, Sharad Kumar). Corresponding authors: Sharad Kumar, Yoon Lim.

## Abstract

Mitotic catastrophe (MC) is an important oncosuppressive mechanism that serves to eliminate cells that become polyploid or aneuploidy due to aberrant mitosis. Previous studies have demonstrated that the activation and catalytic function of caspase-2 are key steps in MC to trigger apoptosis and/or cell cycle arrest of mitotically defective cells. However, the molecular mechanisms that regulate caspase-2 activation and its function are unclear. Here we identify six new phosphorylation sites in caspase-2 and show that a key mitotic kinase, Aurora B kinase (AURKB), phosphorylates caspase-2 at the highly conserved residue S384. We demonstrate that phosphorylation at S384 blocks caspase-2 catalytic activity and apoptosis function in response to mitotic insults, without affecting caspase-2 dimerisation. Moreover, molecular modelling suggests that phosphorylation at S384 may affect substrate binding by caspase-2. We propose that caspase-2 S384 phosphorylation by AURKB is a key mechanism that controls caspase-2 activation during mitosis.

## Introduction

Mitosis is a highly dynamic process that is tightly regulated by a series of surveilling mechanisms that ensure the faithful segregation of chromosomes into the nascent daughter cells, thereby maintaining genomic integrity. Errors in properly partitioning the chromosomes (karyokinesis) and/or separating the two daughter cells (cytokinesis) during cell division generate cells carrying abnormal genomic contents (i.e., aneuploidy or polyploidy) ^1^. The chromosomal passenger complex (CPC) is a key factor in controlling the proper execution of both karyokinesis and cytokinesis, especially through the activity of its kinase subunit Aurora B (AURKB), to prevent the formation of aneuploid and polyploid cells^2^. However, when mitotic errors cannot be corrected, they ultimately trigger mitotic catastrophe (MC), a key mechanism that prevents the proliferation and survival of mitotically aberrant cells through either regulated cell death (RCD), such as apoptosis, or cellular senescence ^1, 3^. Failure of the MC process leads to persistent genomic instability, an enabling characteristic and hallmark of cancer^4^. Therefore, MC is a critical mechanism to prevent cells from becoming tumourigenic.

Caspase-2, the most evolutionarily conserved member of the mammalian caspase family, was discovered over two decades ago^5^. Previous studies using knockout (KO) mice have demonstrated that caspase-2 deficiency promotes tumour development following replicative or oncogenic stress ^6, 7, 8, 9, 10^. Recent studies have demonstrated that caspase-2 activation and its catalytic activity are critical steps in MC signalling. Our previous studies demonstrated that caspase-2-deficient cells acquire extensive aneuploidy following replicative stress in culture ^11^, following prolonged mitotic arrest caused by inhibition of polo-like kinase 1 (Plk1) ^12^ and in the bone marrow of aged caspase-2 KO mice^13^. This was partly caused by decreased Bid cleavage, reduced cell death, and clonogenic survival of aberrant mitotic cells ^12^. Importantly, cells from mice harboring catalytic-dead caspase-2 (*CASP2^C320S^)* also show increased aneuploidy following prolonged mitotic arrest^12^. These findings suggest that the activation and enzymatic activity of caspase-2 are required to mediate apoptosis of aneuploid cells. Other studies have demonstrated that caspase-2 activation following cytokinesis failure following AURKB inhibition triggers cleavage of MDM2, leading to p53 stabilisation and cell cycle arrest, as an alternative mechanism to prevent aneuploidy and polyploidy^14, 15^. Although this evidence shows that caspase-2 activation and its activity are required to prevent survival and proliferation of cells with mitotic defects, the molecular mechanisms have not been well defined.

Previous studies indicate that activation of caspase-2 can be modulated by phosphorylation at various sites to inhibit its activation in different experimental conditions^16, 17, 18^. Phosphorylation at S308 (S340 in human) in *Xenopus* caspase-2 during mitosis is mediated by Cdk1-cyclin B1 to protect cells from incidental apoptotic cell death. Under these conditions, caspase-2 activation levels are balanced by protein phosphatase 1 (PP1)-mediated S308 dephosphorylation^16^. Furthermore, in *Xenopus* oocytes, S135 (S164 in human) of caspase-2 is phosphorylated under nutrient rich conditions, by calcium/calmodulin-dependent kinase II (CaMKII) to prevent its activation^18^. Phosphorylation at S157 by casein kinase-2 (CK2) also inhibits activation of caspase-2 in TNF-alpha-related apoptosis-inducing ligand (TRAIL)-mediated apoptosis ^17^. Interestingly, residues S157 and S164 are positioned within the linker region between the caspase activation and recruitment domain (CARD) and the large (p19) subunit, and residue S340 maps between the large (p19) and small (p14) subunits of caspase-2, suggesting that phosphorylation at these sites acts to prevent caspase-2 processing and activation. Recently, it has also been shown that phosphorylation at T180 of caspase-2 by mitogen-activated protein kinase (MAPK), is involved in the sterol regulatory element-binding protein (SREBP) activation process to regulate lipid metabolism ^19^. However, it is not known whether phosphorylation at these sites regulates caspase-2 activation and its activity in response to aberrant and/or failed mitosis.

In this study, we identified a number of previously unknown phosphorylation sites in caspase-2. Specifically, we demonstrate AURKB phosphorylates caspase-2 at the highly conserved S384 residue within the small subunit. Furthermore, our data indicate an alternative MC regulatory mechanism through AURKB-mediated inhibitory phosphorylation of caspase-2 and suggest inhibition of AURKB activity is required to trigger apoptosis or cell cycle arrest following failed mitosis.

## Results

### Identification of phosphorylation sites in caspase-2

Previous studies have shown that phosphorylation can inhibit caspase-2 activation or regulate its interaction with other molecules under different physiological conditions ^16, 17, 18, 19^. Therefore, we set out to define all potential caspase-2 phosphorylation sites in viable cells in culture. To identify phosphorylation sites in caspase-2, LC-MS/MS experiments were conducted using trypsin-digested GFP immunoprecipitates from U2OS-*CASP2^-/-^* cells transiently expressing GFP-tagged catalytically-inactive mouse caspase-2-C320G. (Fig. 1a and Supplementary Fig. 1). Supplementary Fig. 2 shows the mass spectra used for high-confidence identification of caspase-2 phosphorylation sites. The annotated peptide sequence demonstrates all of the peptide fragments used for phosphorylation site identification. In total we identified 11 phosphorylation sites in caspase-2 (Fig. 1b and 1c), including previously characterised residues S157, S164, T180 and S340, ^16, 18, 19^, and 6 novel sites S24, S80, T161, S220, S346, S384. Phosphorylation at T158 in human caspase-2 has also been reported in the PHOSIDA database, but its role is functionally undefined ^20^.

**Figure 1.**
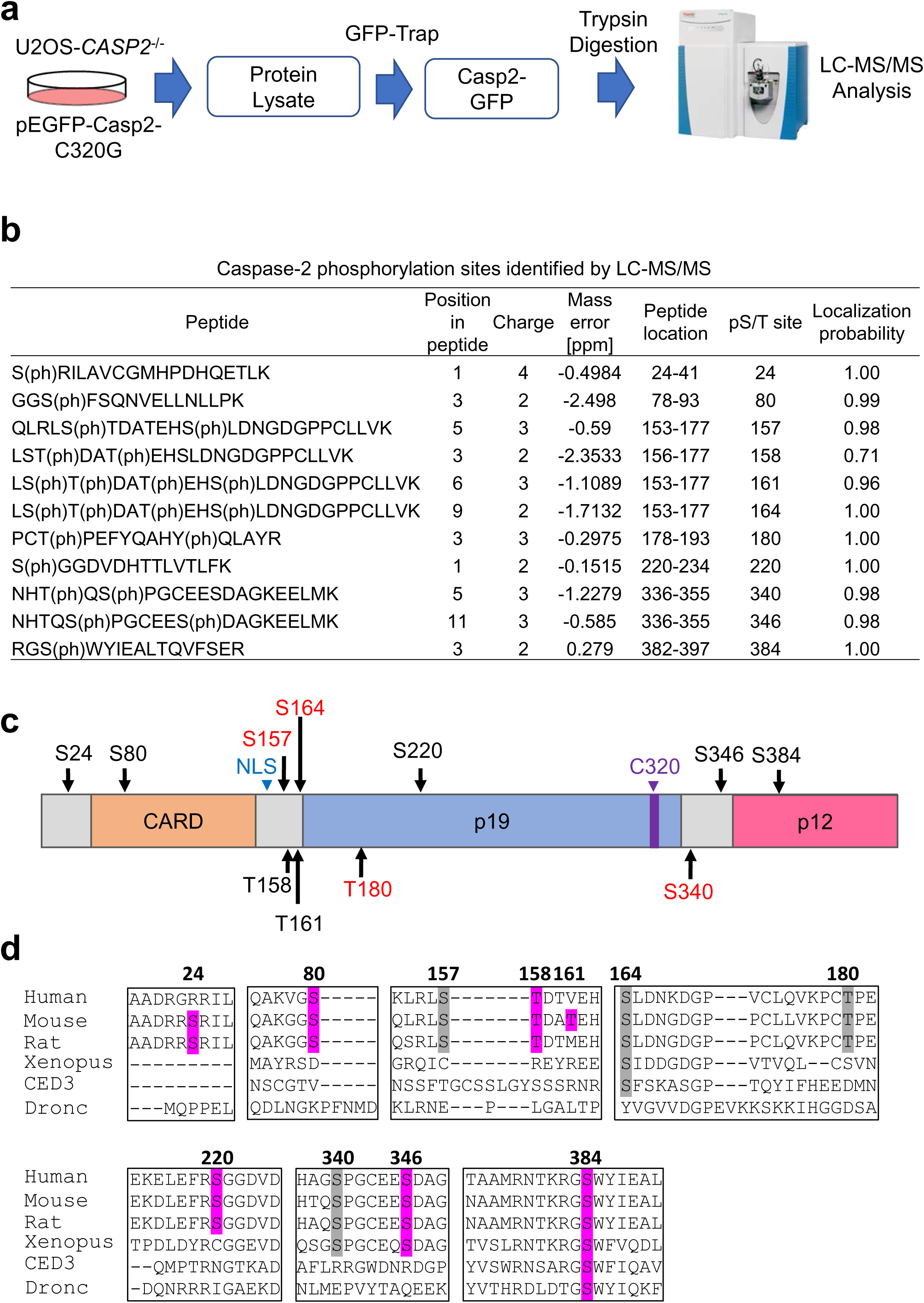
Proteomic analysis identified six new phosphorylation sites in caspase-2. **a.** Schematic diagram of phosphorylation site analysis of caspase-2. **b.** Caspase-2 phosphorylation sites identified by LC-MS/MS. The phosphorylated amino acid position, charge, mass error, peptide location in caspase-2, phosphorylated S/T site position in caspase-2 and localisation probability are indicated. Phosphorylated residues are indicated in brackets. Spectra for the peptides in Table 1 can be found in Supplementary Fig. S2. **c.** Phosphorylation sites in caspase-2 identified by mass spectrometry. Functionally reported sites are indicated in red. S (Ser) or T (Thr) indicates positions of novel potential phosphorylation sites. NLS, nuclear localisation signal. C320, catalytic Cys. **d.** Multiple amino acid sequence alignment showing conservation of phosphorylation sites in caspase-2 from different species. Newly identified and known phosphorylation sites with high homology are shaded in purple and grey respectively.

Next, we analysed the evolutionary conservation of the identified phosphorylation sites and found that S80, S157, T158, T180, S220 are conserved in mammals whereas S340 and S346 are also conserved in *Xenopus* caspase-2. Interestingly, S164 and S384 are most highly conserved, especially S384, which is remakably well conserved among caspase-2 in across the different species including in *Drosophila* apical caspase Dronc and *C. elegans* CED3 (Fig. 1d). It is important to note that Dronc and CED3 respectively are the only CARD containing caspases in flies and nematodes and are functionally analogous to both mammalian caspase-2 and caspase-9^21, 22^. This finding suggests that caspase-2 S384 residue might be functionally important.

### Phosphorylation regulates caspase-2 activation and function

Previous studies have demonstrated that the first step in caspase-2 activation is homo-dimerisation via its CARD, followed by auto-processing ^23, 24, 25^. To functionally characterise the newly identified phosphorylation sites in caspase-2, we generated phospho-mimetic (Ser/Glu) and phospho-deficient (Ser/Ala or Thr/Val) mutants for each residue and then examined whether these mutants affected the processing and activation of caspase-2, by assessing the cleavage of its substrates Bid and MDM2 ^26, 14, 15^. We transiently expressed GFP-tagged caspase-2 phosphorylation site mutants, WT or the catalytically-inactive mutant C320G, in U2OS-*CASP2^-/-^* cells that we had generated using CRISPR/Cas9. As expected, WT caspase-2 but not the C320G mutant cleaved MDM2, generating a p60 cleavage fragment (60 kDa) (Fig. 2a). We also found that most of the phosphorylation mutants could cleave MDM2 to some extent. In contrast, MDM2 cleavage was completely abolished by expression of the phosphomimetic S384E caspase-2 mutant (Fig. 2a). To further confirm cleavage of Bid, we co-expressed Bid-HA with the WT and mutant caspase-2 constructs and monitored loss of the full-length Bid protein. Similarly, we found that all the phosphorylation site mutants except S384E showed reduced levels or loss of full-length Bid, indicating complete cleavage. As expected, the expression of the C320G mutant did not induce cleavage of Bid (Fig. 2b). We also assessed the processing of the transfected procaspase-2 mutants and observed reduced cleavage of S384E, to a similar extent as seen with C320G, as indicated by higher levels of full-length caspase-2-GFP protein (74 kDa) and reduced levels of the 18 kDa subunit band compared to WT caspase-2 or other phosphosite mutants (Fig. 2a and b). These results demonstrate that the caspase-2 phospho-mimetic S384E mutation affects its auto-processing and catalytic activity. Consistent with these findings, and similar to the C320G mutant, the expression of S384E also exhibited reduced ability to promote cell death compared to WT or S384A in *Casp2^-/-^* immortalised mouse embryonic fibroblasts (MEFs) (Fig. 2c and d)

**Figure 2.**
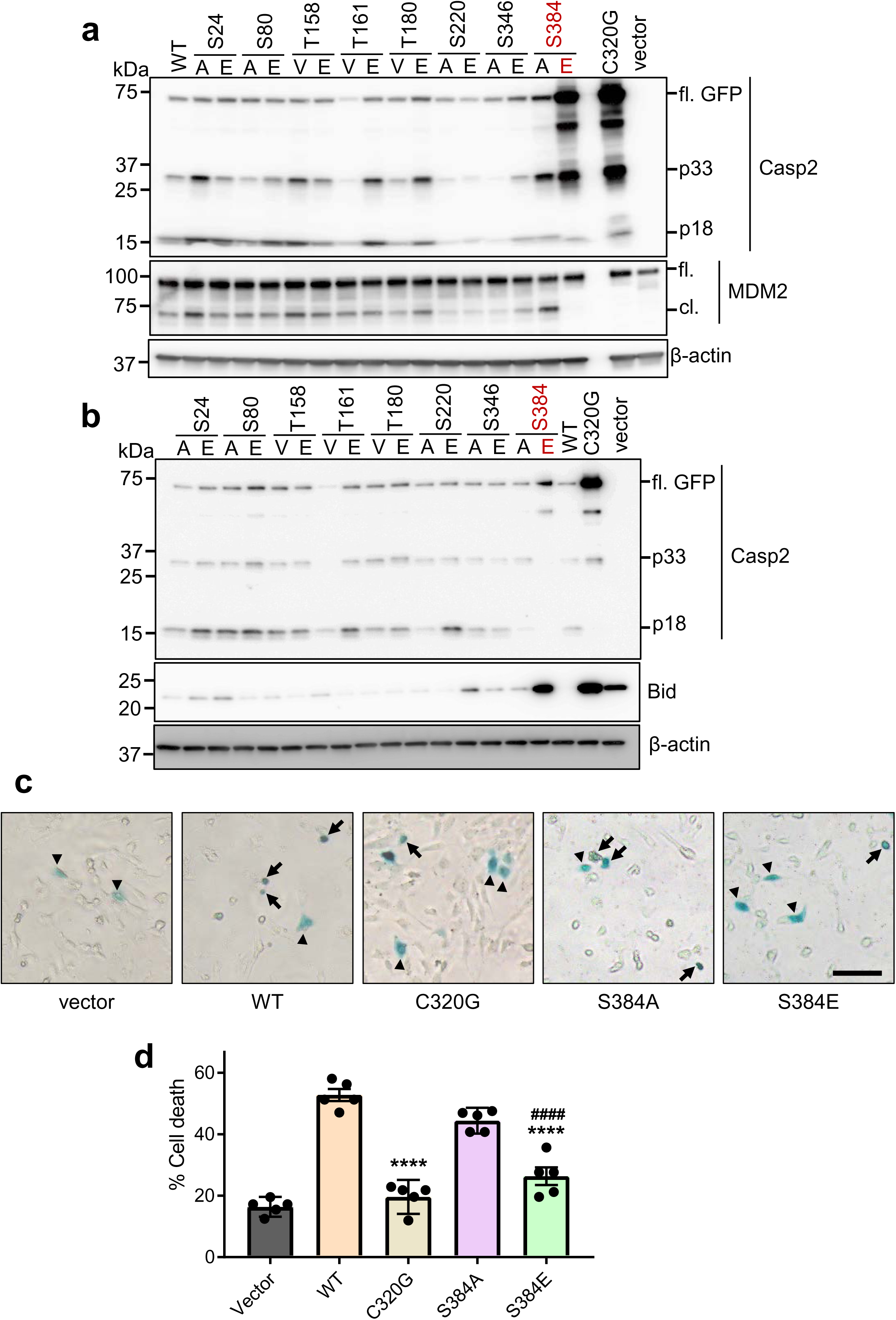
Caspase-2-S384E mutant does not cleave MDM2 or Bid. **a.** U2OS-*CASP2*^-/-^ cells were transfected with GFP mock vector, GFP-caspase-2 WT, C320G or phosphorylation site mutants. Cell lysates were subjected to immunoblotting. with the indicated antibodies. β-actin was used as loading control. **b.** U2OS-*CASP2*^-/-^ cells were co-transfected with GFP mock vector, GFP-caspase-2 WT, C320G or phosphorylation site mutants and with Bid-HA. Cell lysates were subjected to immunoblotting with the indicated antibodies. β-actin was used as loading control**. c and d.** For cell death assays, *Casp2^-/-^* immortalised MEFs were co-transfected with GFP-caspase-2 WT, C320G, S384A or S384E and β-gal expressing plasmids. After 24 h, cells were fixed and incubated with an X-gal containing solution. Blue (transfected) cells were counted for apoptotic morphology under microscope. **c.** Rrepresentative images showing X-gal staining with live or apoptotic cell morphology. Arrow, dead cell; arrowhead, live cell. Scale bar=50μm **d**. Graph showing % cell death. *, vs. WT; ****, p<0.0001; ^#^, S384A vs. S384E; ^####^, p<0.0001; mean ± SEM. One-way ANOVA with post hoc test.

We next assessed the enzymatic activity of the various caspase-2 phosphorylation mutant cell lysates using an *in vitro* caspase activity assay with the selective fluorogenic caspase-2 substrate, VDVAD-AFC ^27, 28^. Interestingly, several mutants showed significantly reduced VDVADase cleavage activity, including T180V, S220A, S220E, S346A and S384A (Supplementary Fig. 3a). Consistent with the caspase-2 processing and substrate cleavage data (Fig. 2), S384E showed no enzymatic activity, similar to C320G or the mock vector control (Supplementary Fig. 3a). As the S384A mutation also showed reduced enzymatic activity, we wanted to confirm that the S384E mutation was indeed acting as a phospho-mimetic and not simply disrupting protein structure and/or folding. Therefore, we generated Ser to Gly (small amino acid) and Ser to Thr (uncharged, polar amino acid like serine) mutations at this site and assessed MDM2 cleavage following expression of these mutants in U2OS-*CASP2^-/-^* cells. We observed that, like S384A, both the S384G and S384T mutants were still able to cleave MDM2 and could also be processed in transfected cells (Supplementary Fig. 3b), which indicates that the loss of substrate cleavage ability by the S384E mutant is not caused by random amino acid substitution that may influence protein conformation at this site. Together, these results indicate that phosphorylation at S384 inhibits caspase-2 activation and activity in cells and that several phosphorylation site mutations can affect the enzymatic activity of caspase-2, but they do not impair its substrate cleavage and cell death inducing activity.

### Caspase-2-S384E does not disrupt dimerisation

As homodimerisation is sufficient for initial caspase-2 activation^23^, we examined whether the S384E mutation affected caspase-2 dimerisation using the bimolecular fluorescence complementation (BiFC) assay, a visualizing tool of protein-protein interaction^12, 29^, with full-length caspase-2-S384E and CASP2-C320A. We blocked mitosis using the Plk1 inhibitor BI2536 (BI), which we have previously demonstrated to result in caspase-2 dimerisation^12^. There were no differences in dimerisation of caspase-2-S384E or CASP2-C320A between control (DMSO) and BI treatment (Fig. 3a and 3b). We also expressed GST recombinant caspase-2-WT, C320G or S384E in *E.coli* and found that WT GST-caspase-2 was cleaved to form an intermediate (∼50kD) and fully processed large subunit (18kDa), whereas GST-caspase-2-S384E was not processed and behaved like the recombinant C320G protein (Fig. 3c). This demonstrates that caspase-2 S384E substitution prevents autoprocessing of caspase-2. Caspase assays using VDVAD-AMC showed robust catalytic activity by WT caspase-2 but not by both the C320G and S384E mutants (Fig. 3d), which further confirmed that caspase-2-S384E mutant protein lacks catalytic activity. Our data suggest that phosphorylation at S384 does not prevent the initial caspase-2 dimerisation event, but inhibits autoproteolysis and downstream activation, and therefore likely exerts a dominant negative function.

**Figure 3.**
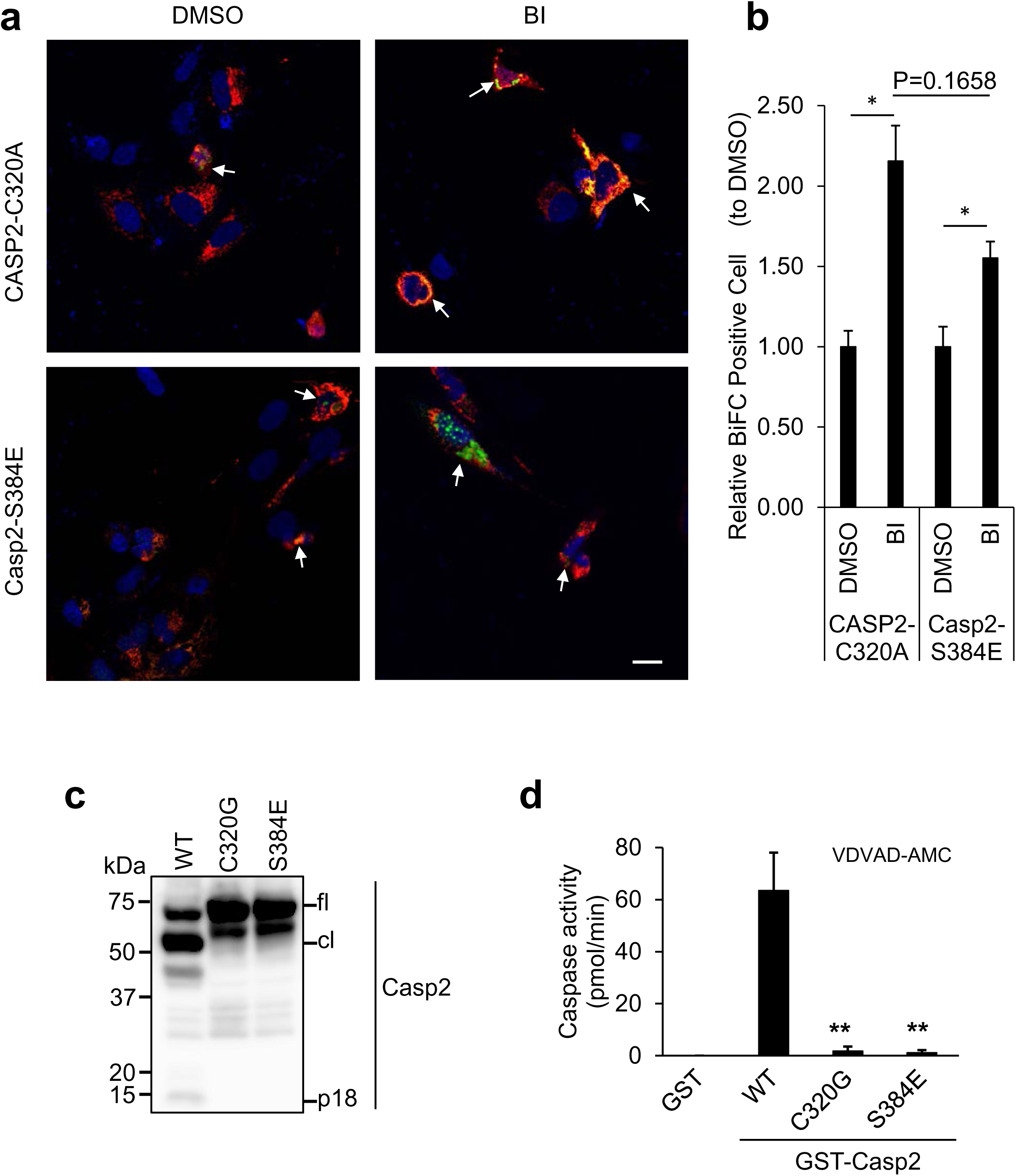
Caspase-2-S384E mutant can still homo-dimerise but lacks enzymatic activity. **a.** Representative confocal images showing dimerisation of Casp2-S384E or CASP2-C320A by BiFC. U2OS-*CASP2^-/-^* cells were transfected with Casp2-S384E or CASP2-C320A BiFC constructs, respectively, and treated with DMSO or 100 nM BI2536 (BI) for 24h. All cells were also treated with zVAD (20 µM), pan-caspase inhibitor. Arrows indicate dimerised caspase-2. Scale bar=20μm. **b.** Quantitation of relative BiFC positive cells are indicated. *, P<0.05. **c.** Immunoblot analysis recombinant GST-caspase-2-WT, C320G and S384E protein expression and auto-processing. **d.** Caspase activity of recombinant GST-caspase-2-WT, C320G and S384E protein, was assessed by cleavage of VDVAD-AMC. Data represented as mean ± SEM from three independent experiments. *, vs. WT; **, p<0.01.

### Caspase-2 S384 is a target of Aurora B kinase

To identify potential kinases that may phosphorylate S384 in caspase-2, we used various computational tools and databases including PHOSIDA, Scansite and KinasePhos 2.0. These analyses suggested that S384 might be a target site for AURKB. The consensus phosphorylation motifs for AURKB are (K/R)1-3-X-(S/T) or (K/R)-(R/K)-X0-2-(S/T) where X indicates any amino acid^30, 31^. Such a sequence is present around S384 in caspase-2 (Fig. 1d). Therefore we first performed a GST-Casp2 pull-down assay to examine whether AURKB directly binds to caspase-2. Recombinant GST or GST-Casp2-C302G (Casp2-FL) proteins were incubated with purified His-tagged AURKB and we found that GST-Casp2 proteins coprecipitated with AURKB whereas GST alone did not (Fig. 4a). Next, we performed an *in vitro* phosphorylation assay using AURKB with either GST-Casp2 full length (Casp2-fl) WT or the S384A mutant as substrates. The result demonstrated that full-length caspase-2 is phosphorylated by AURKB (Fig. 4b). To further validate phosphorylation at S384 by AURKB, we used a small peptide fragment of caspase-2 comprising S384 (aa 363–423; Casp2_363-423_; Fig. 4b). This confirmed that S384 of caspase-2 is phosphorylated at S384 by AURKB *in vitro.* Notably, AURKB phosphorylation was almost completely abolished in the Casp2_363-423_ S384A mutant fragment, but was only diminished in the Casp2-fl S384A mutant (Fig. 4b), suggesting that AURKB might phosphorylate additional residues besides S384. Interestingly, AURKA did not phosphorylate S384 (Supplementary Fig. 4), indicating that this residue is a specific site for AURKB.

**Figure 4.**
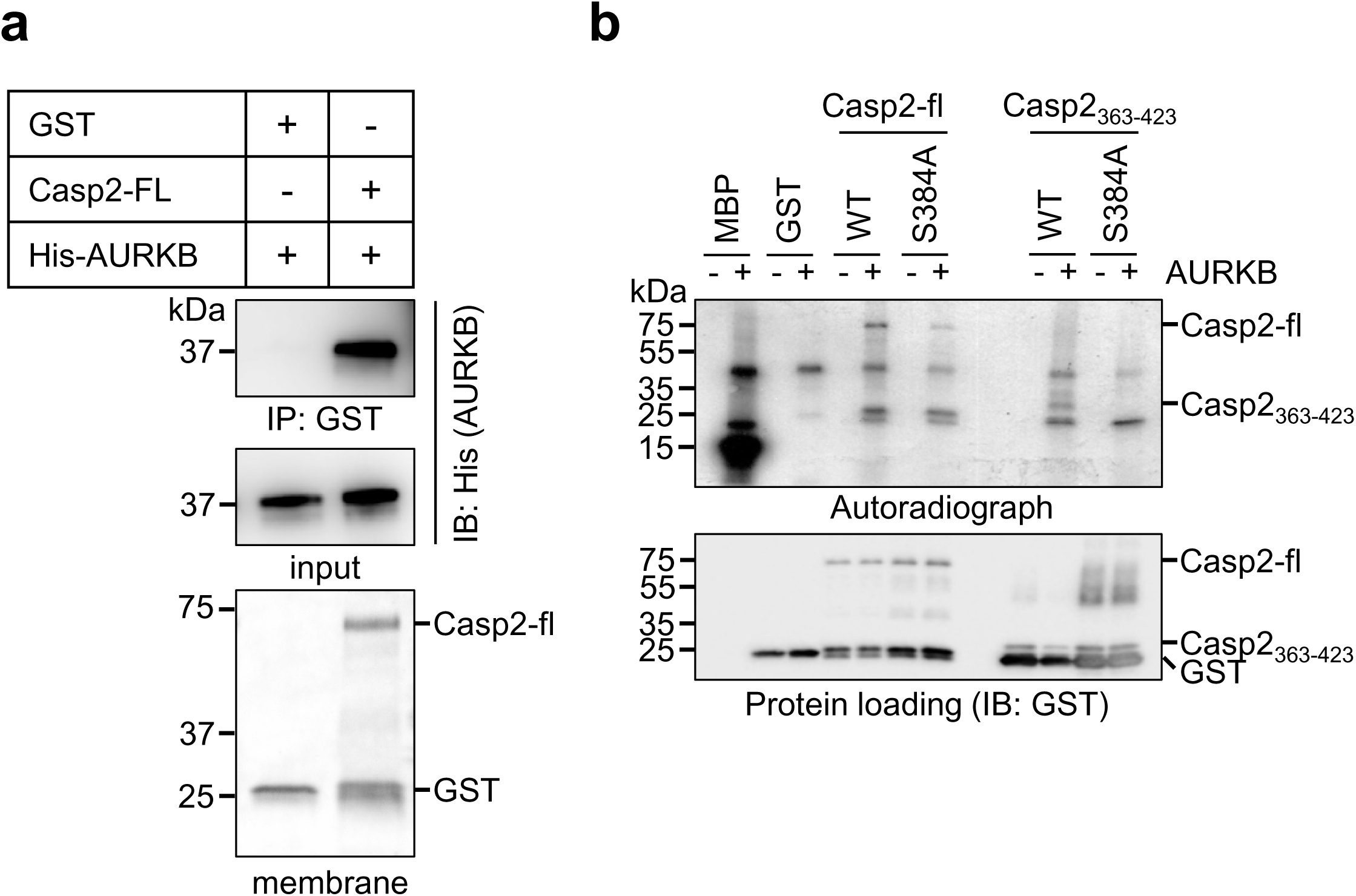
Aurora B kinase phosphorylates caspase-2 at S384 *in vitro*. **a.** *In vitro* interaction between purified recombinant GST-Casp2-C320G (Casp2-FL) and His-AURKB was determined by GST pull-down assay. Stain-free membrane shows protein loading amount for GST or GST-Casp2. Antibodies used are indicated. **b.** GST-Casp2-C320G (S384 WT), GST-Casp2-C320G-S384A (S384A), GST-Casp2_363-423_-WT, GST-Casp2_363-423_-S384A or GST was subjected to *in vitro* phosphorylation by incubation with AURKB and [γ-^32^P]-ATP and analysed by autoradiography. IB with GST antibody shows equal loading. MBP, myeloid basic protein was used as positive control for AURKB phosphorylation.

### Caspase-2-S384E expression results in increased polyploidy and resistance to cell death following mitotic stress

To further characterize the functional consequence of phosphorylation at S384, we established a series of U2OS cells stably expressing GFP (sWT) or U2OS-*CASP2^-/-^* cells stably expressing GFP (sKO), GFP tagged caspase-2-C320G (sC320G) or S384E (sS384E clones #1 and #2) (Supplementary Fig. 5). These cell lines were treated with two drugs, the AURKB inhibitor ZM447439 (ZM) ^32^ and the myosin II inhibitor blebbistatin^33^, for 48 hours to induce polyploidy and then cleavage of caspase-2 and its substrates MDM2 was analysed by immunoblotting and DNA content was measured by flow cytometry. Consistent with previous reports ^14, 15^, sWT cells showed cleavage of caspase-2 (p18 kDa), MDM2 and increased p53 and p21 accumulation in response to AURKB inhibition (Fig. 5a). The sKO cells still showed increased p53 levels following ZM treatment, albeit less than sWT cells, but p21 levels were not increased and cleavage of MDM2 was absent, which is consistent with previously reported findings (Fig. 5a) ^14^. Interestingly, similar to sKO and sC320G, the two independent sS384E cell lines also showed absence of MDM2 cleavage and p21 increase in response to ZM, while still exhibiting induction of p53 (Fig. 5a). DNA content analysis of sKO cells showed significantly increased polyploid (>4n) cell accumulation compared to sWT cells (Fig. 5b and c), which is consistent with previous reports ^14, 15^. Similarly, sC320G and sS384E cell lines also showed increased polyploidy following ZM treatment (Fig. 5b and c).

**Figure 5.**
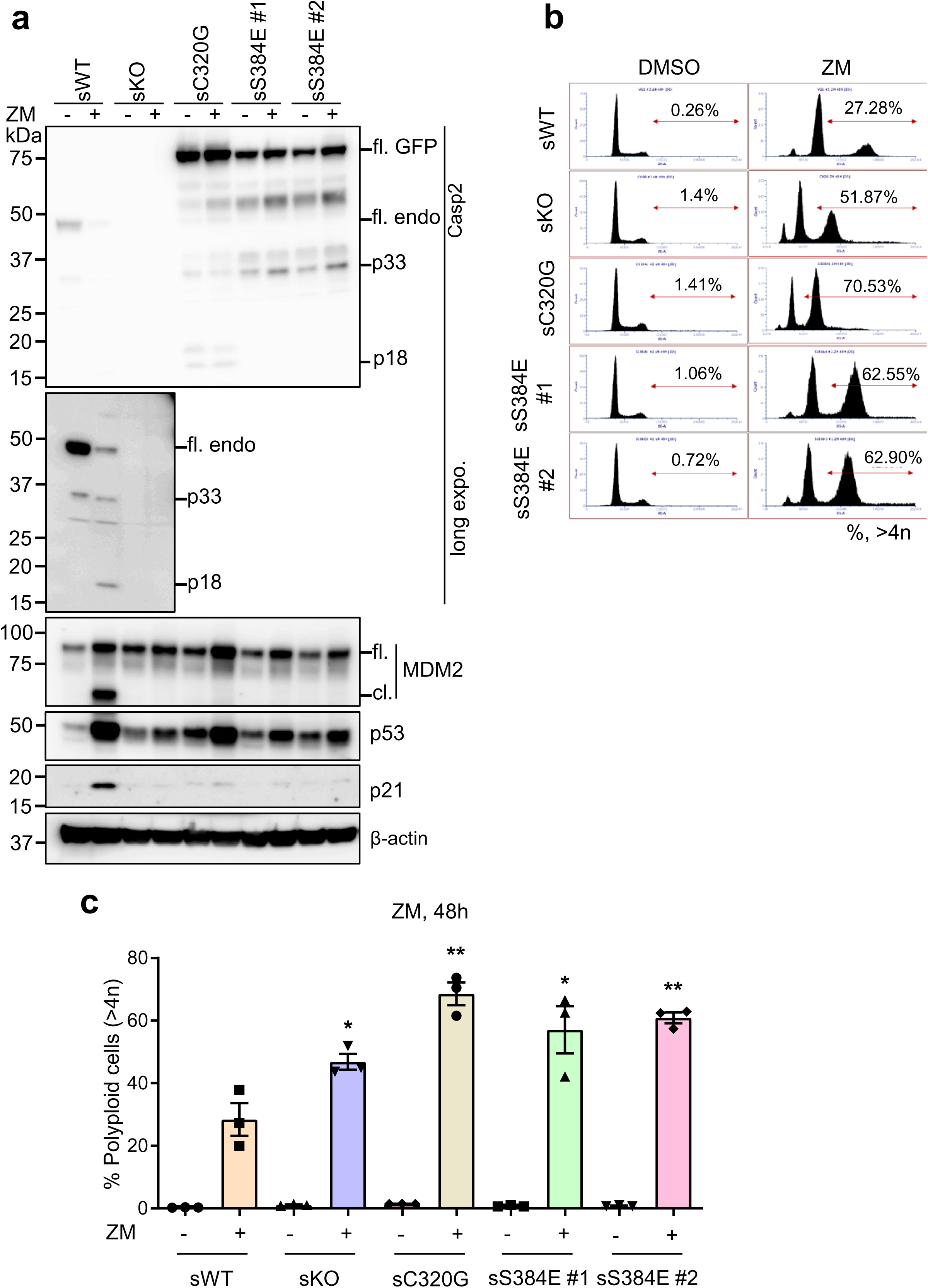
Caspase-2-S384E expressing cells fail to cleave MDM2 and show increased polyploidy following AURBK inhibition. GFP-expressing U2OS (sWT) and U2OS-*CASP2^-/-^* expressing GFP (sKO), GFP-caspase-2-C320G (sC320G) or GFP-caspase-2-S384E (sS384 #1 and #2) were treated with DMSO or 2 µM ZM447439 (ZM) for 48 h and subjected to immunoblot and DNA content analysis. **a.** Representative immunoblots (of three independent experiments performed) of cell lysates from treated stable cell lines. Antibodies used for immunoblotting are as indicated. β-actin was used as loading control. **b.** Representative flow cytometric profiles of the DNA content in cells following ZM treatment. Percentage of polyploid cells (> 4n) is indicated. **c.** Graph comparing percentage of cells with polyploid (>4N) DNA content following ZM treatment. mean ± SEM; n=3. *, vs. WT + ZM; *, p<0.05; **, p<0.01.

We used also an alternative approach to induce polyploidy in by treating them with blebbistatin to inhibit non-muscle myosin II, which plays an essential role in the constriction of the actomyosin contractile ring during cytokinesis ^33, 34^. Importantly, similar results were observed when the U2OS stable cell lines were treated with blebbistatin, although the accumulation of p53 reduced (Fig. 6a), particularly when compared to treatment with the AURKB inhibitor, AZD1152 ^32^ (Supplementary Fig. S6a) in the U2OS parental and *CASP2^-/-^* cells. The percentage of cells with polyploidy was also significantly increased in sKO, sC320G and sS384E cell lines compared to the sWT although the increase is not as robust as AURKB-I treatment (Fig. 6b and Supplementary Fig. S6b).

**Figure 6.**
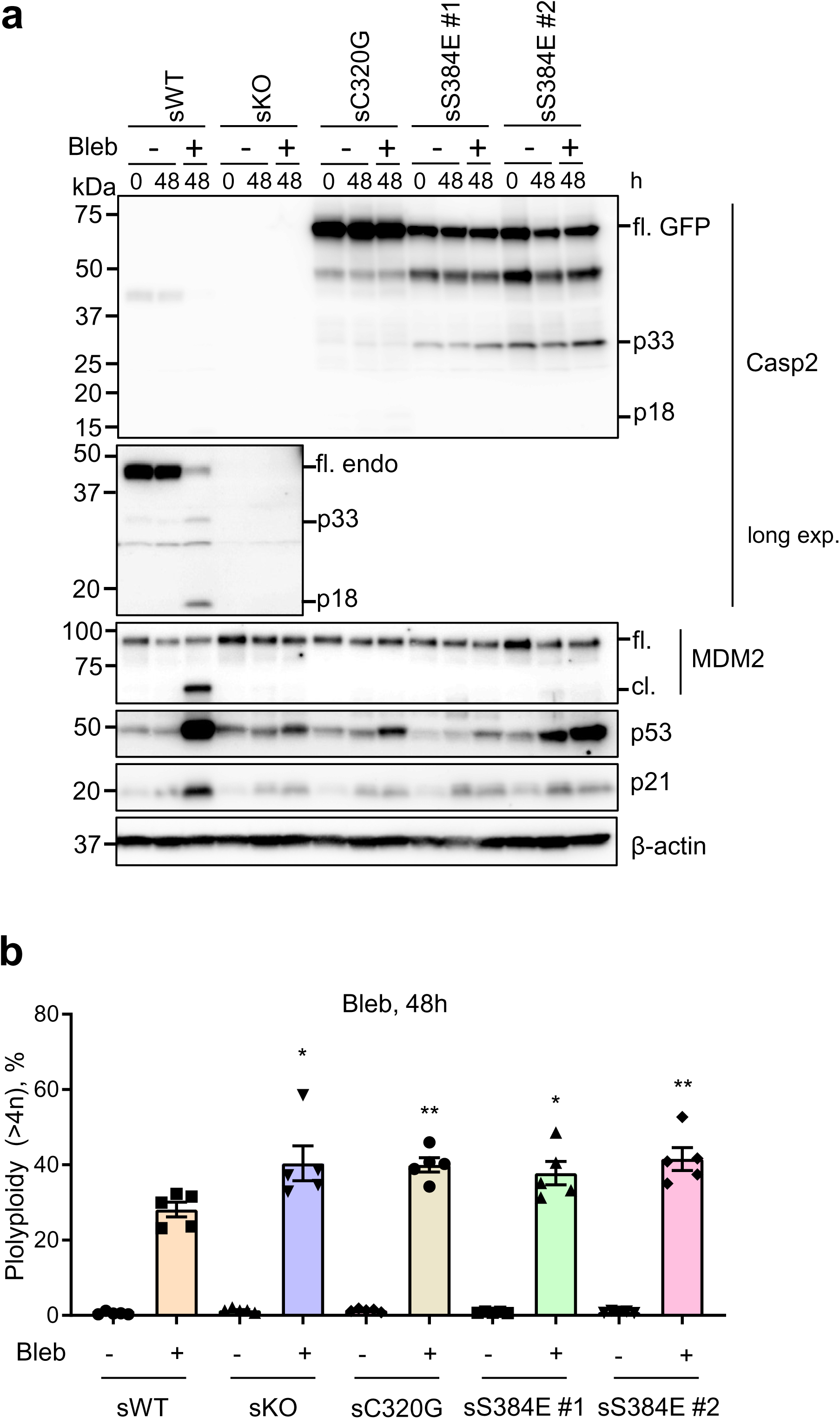
Caspase-2-S384E expressing cells fail to cleave MDM2 and show increased polyploidy following cytokinesis block with blebbistatin. GFP-expressing U2OS (sWT) and U2OS-*CASP2^-/-^* expressing GFP (sKO), GFP-caspase-2-C320G (sC320G) or GFP-caspase-2-S384E (sS384 #1 and #2), were treated with DMSO or 50 µM blebbistatin (Bleb) for 48 h and subjected to immunoblot and DNA content analysis. **a.** Representative immunoblots (of five independent experiments performed) of cell lysates from treated stable cell lines. Antibodies used for immunoblotting are as indicated. β-actin was used as loading control. **b.** Graph comparing percentage of cells with polyploid (>4N) DNA content following Bleb treatment. mean ± SEM; n=5. *, vs. WT + Bleb; *, p<0.05; **, p<0.01.

To examine the effect of S384E following different mitotic stresses, we also treated the U2OS stable cell lines with the Plk1 inhibitor BI2536 (BI), which induces mitotic arrest, chromosome missegregation, aneuploidy and caspase-2 dependent apoptotic cell death ^12^. BI-treated cells were subjected to cell viability assay and assessed for Bid cleavage and apoptosis. Following treatment with BI for 48 h, sKO, sC320G and sS384E cell lines were more resistant to apoptosis than sWT cells (Fig. 7a). The sWT cells also showed caspase-2 activation, cleavage of PARP, caspase-3 and Bid, demonstrating BI2 induction of cell death (Fig 7b). As previously reported, sKO and sC320G cells showed reduced cleavage of PARP, Bid and caspase-3 ^12^ and this was also observed in the sS384E cell lines (Fig. 7b). Taken together, these experiments confirmed that phosphorylation at S384 in caspase-2 inhibits caspase-2 activity and cell death function.

**Figure 7.**
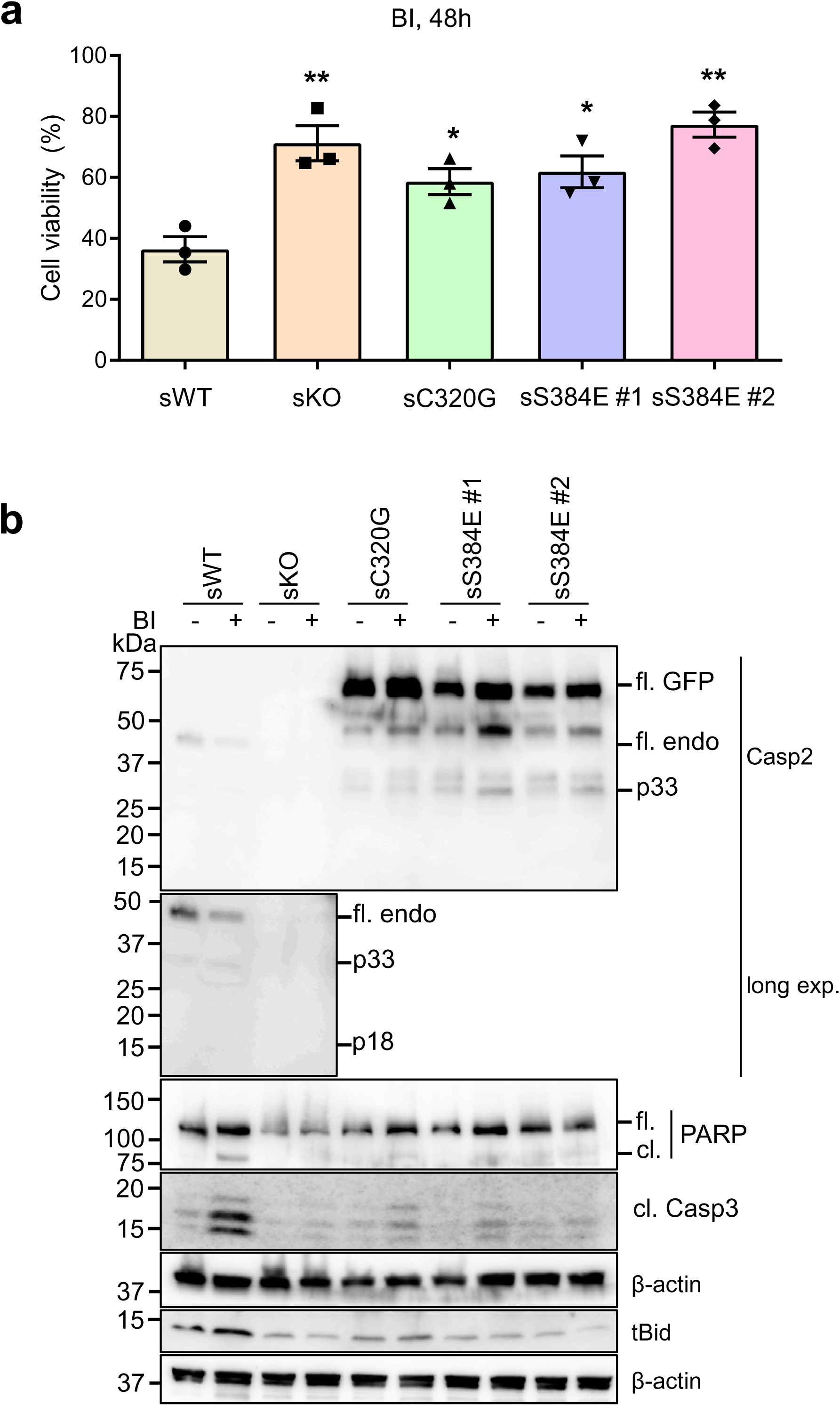
Caspase-2-S384E expressing cells are resistant to aberrant mitosis-mediated cell death. GFP-expressing U2OS (sWT) and U2OS-*CASP2^-/-^* expressing GFP (sKO), GFP-caspase-2-C320G (sC320G) or GFP-caspase-2-S384E (sS384 #1 and #2), were treated with DMSO or 100 nM BI 2536 (BI) for 48 h and subjected to immunoblot and MTS assay. **a.** Plot showing percentage of viable cells following BI treatment. mean ± SEM; n=3. *, vs. WT; *, p<0.05; **, p<0.01. **b.** Representative immunoblots (of three independent experiments performed) of cell lysates from treated stable cell lines. Antibodies used for immunoblotting are as indicated. β-actin was used as loading control.

## Discussion

In the present study, we identified six new phosphorylation sites in mouse caspase-2 by LC-MS/MS. We used the catalytically-inactive caspase-2 C320G mutant, because unlike WT caspase-2, it is not associated with cytotoxicity when expressed ectopically^23^. This allowed us to identify new phosphorylation sites associated with full length caspase-2 zymogen. Of the six newly identified phosphorylation sites we found that the S384, which is highly conserved, is critical for the regulation of caspase-2 activity and function. Our studies utilising reexpression of caspase-2 mutants in caspase-2 deficient cells demonstrated that the caspase-2 S384E phospho-mimetic mutant prevents caspase-2 activation and its catalytic activity. First, S384E caspase-2 shows reduced autoprocessing of recombinant protein in cells. While we observed some caspase-2 cleaved products (p33 and p18), in S384E and C320G overexpressing cells, recombinant protein did not show caspase-2 catalytic activity, indicating this cleavage is likely a caused by other cellular proteases or caspase-3^35^. Secondly, S384E was unable to cleave the known caspase-2 substrates, MDM2 and Bid. Thirdly, caspase-2-S384E can still homodimerise but is catalytically inactive. Lastly, expression of the caspase-2-S384E shows predominantly nuclear localisation, similar to the C320G mutant, which may also impede activation by other caspases in the cytosol. Our results suggest that while S384E does not affect dimerisation, the first step of caspase-2 activation, it prevents autoproteolytic processing, cytosolic localisation and complete caspase-2 activation. This may be in part caused by steric hindrance on the substrate binding site. These data suggest that S384E caspase-2 likely acts as a dominant negative protein.

Caspase-2 is the only member of the caspase family that prefers pentapeptide, such as VDVAD or ADVAD, as substrates rather than a tetrapeptide^27, 28, 36^. T380 and Y420 are key residues that have been shown to be critical for binding the P5 residue^37^. These residues forms hydrogen bonds with P5 Val in VDVAD or Ala in ADVAD pentapeptide substrates^37^. Also, W385 forms a hydrogen bond with P4 Asp. Interestingly, S384 localises next to W385 and close to T380 even although it may not directly interact with the substrate^37^. This suggests that phosphorylation at S384 may cause an interference with substrate binding (Fig. 8a and b) preventing the catalytic activity of caspase-2.

**Figure 8.**
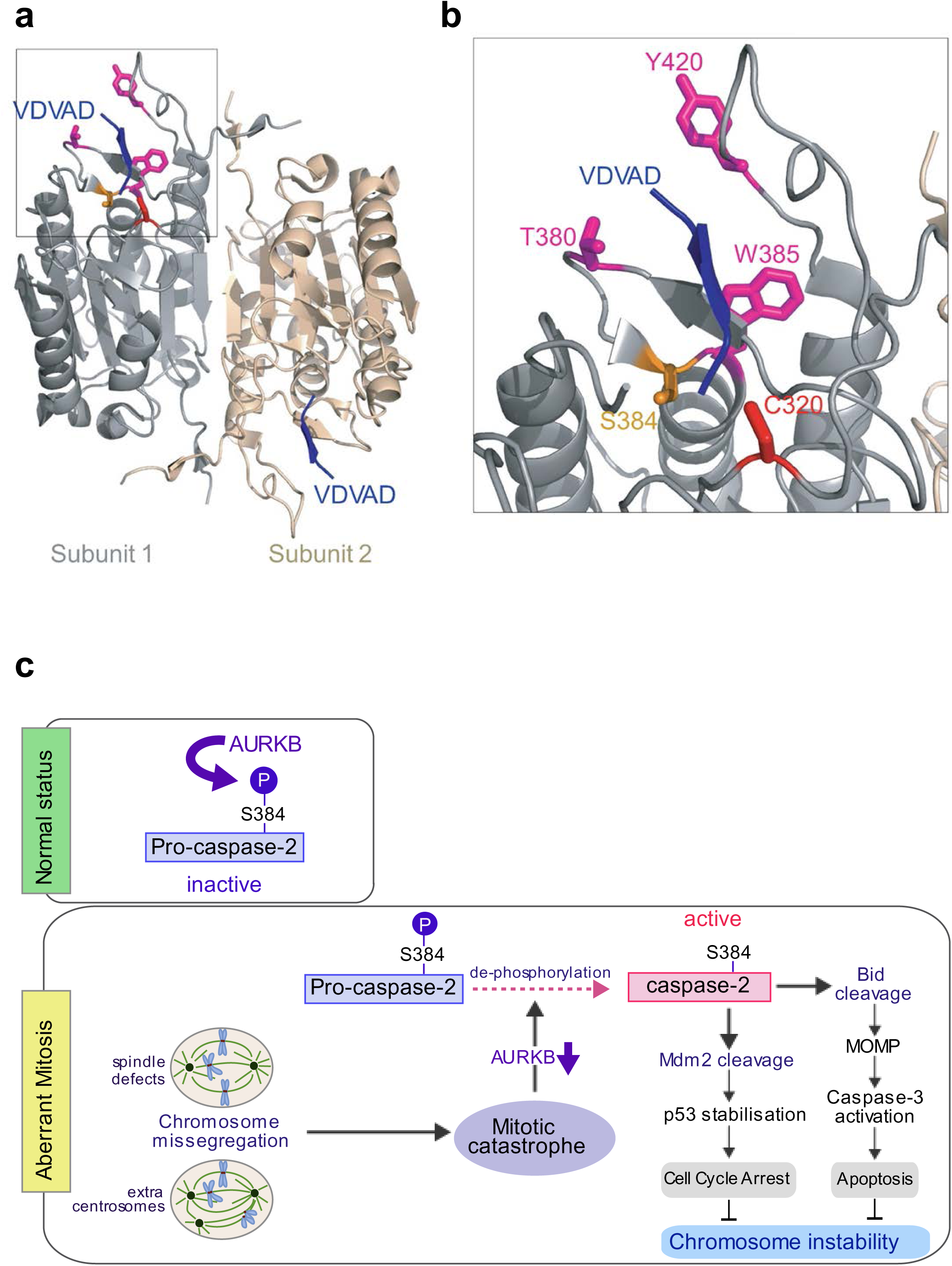
Phosphorylation at S384 affects the conformation of the substrate binding pocket. **a.** The crystal structure of human active caspase-2/VDVAD (PDB ID: 3R6G)^37^ complex. Crystalised caspase-2 is composed of two p19/p12 heterodimers. **b.** Magnified view showing position of S384 and surrounding amino acid residues important for substrate binding. T380 and Y420 and W385 are shown in magenta and interact with P5 and P4 residues^37^. S384 and C320 are shown in orange and red, respectively. **c.** Schematic showing regulation of caspase-2 by AURKB-mediated phosphorylation in aberrant mitosis. In normal status, S384 of caspae-2 is phosphorylated by AURKB, maintaining caspase-2 inactive. In response to aberrant mitosis, such as cytokinesis failure and chromosome missegregation, AURKB activity is reduced, likely allowing S384 dephosphorylation by protein phosphatase(s) (PP). This leads to caspase-2 activation that can result in two outcomes: either cleavage of MDM2 which leads to p53 stabilisation and cell cycle arrest; or cleavage of Bid which leads to outer mitochondrial membrane permeabilization (MOMP), caspase-3 activation and apoptotic cell death. These pathways limit cells becoming aneuploid or polyploid, preventing chromosome instability. Dashed lines indicate unknown mechanisms.

Multiple kinases are responsible for phosphorylation of caspases^38^. As mentioned above, CK2, CaMKII and Cdk1 are known to phosphorylate caspase-2. Our study identified AURKB as an important additional kinase that phosphorylates caspase-2 at S384. We also examined other kinases (including CaMKII, PKA, PAK1, Cdk1, Plk1 and MAPK) but none of them phosphorylated the S384 site in caspase-2 (data not shown). AURKB has various critical functions in mitosis, including chromatin condensation, kinetochore-microtubule error correction, and cytokinesis^39^. Therefore, our findings suggest that AURKB phosphorylation maintains caspase-2 in an inactive state throughout cell division. Notably, caspase-2 is also regulated by Cdk1-mediated inhibitory phosphorylation at S340 during mitosis^16^. However, while Cdk1 activity is inhibited following degradation of its cyclin B partner at anaphase onset, AURKB remains active until late cytokinesis^40^. Importantly, AURKB, along with its CPC partners, can delay the final separation (i.e., abscission) of the two daughter cells in the presence of lagging chromatin at the cleavage site to prevent aneuploidy and polyploidy^41^. Therefore, it appears that caspase-2 is under tight regulation by multiple mitotic kinases to prevent any accidental cell death during cell division. Our studies using the S384E mutant expression in U2OS cells and recombinant S384E caspase-2 protein *in vitro* demonstrate that phosphorylation at S384 prevents caspase-2 catalytic activity and function. Therefore, the inhibition of S384 phosphorylation by targeting AURKB may be an effective mechanism of activating caspase-2 in cycling cells. However, it would be important to decipher whether S384 dephosphorylation could effectively activate caspase-2 during other cell cycle stages or in senescent cells. Interestingly, blebbistatin, which does not inhibit AURKB, also causes cytokinesis failure and caspase-2 activation indicating potentially differential regulation of caspase-2 at different mitotic stages. Importantly, the caspase-2 S384E phospho-mimetic mutant was also not activated following blebbistatin treatment, suggesting an important role for AURKB-mediated phosphorylation in regulating caspase-2 activation during general cytokinesis failure. It has been shown that PP1 can dephosphorylate caspase-2 S308 residue, leading to caspase-2 activation and cell death ^16, 42^. Therefore, identification of the phosphatase(s) of S384 that can trigger caspase-2 activation may provide an important mechanism to eliminate polyploid cells.

In sum, we propose that, in normal mitosis, caspase-2 is maintained in an inactive state by phosphorylation by both Cdk1 and AURKB until anaphase onset and then only by AURKB-mediated phosphorylation at S384 until abscission. If MC is triggered before anaphase, by failure of chromosome alignment for example, inhibition of Cdk1 and AURKB and de-phosphorylation at both S340 and S384 would activate caspase-2 to promote cell death. If instead there are irrecoverable problems specifically during cytokinesis, such as persistent lagging chromatin, AURKB inactivation and dephosphorylation at S384 would lead to caspase-2 activation and apoptosis to prevent survival of polyploid cells (Fig. 8c).

Overexpression and enhanced kinase activity of AURKB in various tumour types are well documented and associated with therapy resistance and low survival rate in various cancers. Therefore the AURKB is considered as a potential therapeutic target in cancer ^43^. Together with our data presented here, this suggests that in tumour cells, AURKB may act to maintain caspase-2 in a phosphorylated inactive state, thereby preventing apoptotic cell death and contributing to therapy resistance. We therefore propose that the phosphorylation state of caspase-2 may predict apoptosis sensitivity and treatment response in cancer.

Cytokinesis failure induced by AURKB inhibition, has previously been shown to induce caspase-2 activation, MDM2 cleavage and accumulation of p53 and cell cycle arrest in both U2OS and A549 cells^14, 15^. We have also previously observed increased polyploidy in U2OS-*CASP2^-/-^* cells, as a consequence of the accumulation of uncleaved, full length MDM2, but still associated with increased p53 levels ^14^. This suggested the presence of an MDM2-independent p53 response following aberrant mitosis, in caspase-2 deficient cells^14^. In this study, we now demonstrate that, similar to *CASP2^-/-^* cells or C320G expression, cells stably expressing the caspase-2 S384E mutant acquire increased polyploidy associated with lack of MDM2 cleavage. Importantly, we have observed this using different mitosis inhibitors, AURKB inhibitors and blebbistatin^33^, that induce polyploidy. These data confirm that our results are a consequence of general cytokinesis failure and phosphorylation of caspase-2 at S384 can prevent the canonical MC response and exacerbate polyploidy.

We have previously demonstrated that caspase-2 activation and its catalytic activity are required for Bid-mediated apoptotic cell death of aneuploid cells ^12^. Our findings here now demonstrate that the caspase-2-S384E cell line does not show Bid cleavage and is more resistant to apoptosis caused by Plk1 inhibition. This further suggests that as well as causing accumulation of MDM2, and dampening p53-mediated cell cycle arrest, the S384E mutant prevents caspase-2-mediated apoptosis. Taken together, our data suggest that AURKB-mediated phosphorylation at S384 of caspase-2 is a critical mechanism that regulates its activation and cellular functions. As activation of caspase-2 is required following MC to prevent survival of polyploid cells following failed mitosis and failure to repair damaged DNA^1, 3^, it will be important to examine in future, whether phosphorylation of caspase-2 at S384 also prevents its tumour suppressor function.

## Materials and Methods

### Antibodies

The following antibodies were used: anti–caspase-2 (clone 11B4); anti-GFP (600-101-215, Rockland); anti-MDM2 (OP46, EMD Millipore); anti–p53 (sc-126, Santa Cruz Biotechnology Inc); anti-p21 (5567430, BD Pharmingen); anti-PARP (9542, Cell Signalling Technology); anti-Bid (2002, Cell Signalling Technology); anti–cleaved caspase-3 (9664, Cell Signalling Technology); anti-HA (H6908, Sigma Aldrich); anti-GST (G7781, Sigma Aldrich); anti-His (ab18184, abcam); anti-β-actin (A1978, Sigma Aldrich).

### Cell culture and transfection

U2OS, U2OS-*CASP2^-/-^* and stable cell lines were maintained in Dulbecco’s Modified Eagles Medium (DMEM, Sigma-Aldrich) supplemented with 10% fetal bovine serum (JHR Biosciences), 0.2 mM L-glutamine (Sigma-Aldrich), 15 mM HEPES (Sigma-Aldrich) and 100 µM penicillin/streptomycin (Sigma-Aldrich) in a humidified incubator at 37 °C with 10% CO_2_. Where indicated, cells were treated with 100 nM BI2536 (Axon Medchem, Netherlands), 2 µM ZM447439 (Sigma-Aldrich), 400 nM AZD1152-HQPA (Sigma-Aldrich), or 50 µM blebbistatin (Selleck Chemicals) for 48 h. MEFs were maintained in DMEM (Sigma-Aldrich) supplemented with 0.2 mM L-glutamine (Sigma-Aldrich), 15mM HEPES (Sigma-Aldrich), 10% fetal bovine serum (JHR Biosciences) and 50 µM β-mercaptoethanol (Sigma-Aldrich), non-essential amino-acid mix (Sigma-Aldrich), 100 µM penicillin/ streptomycin (Sigma-Aldrich). Cells were cultured in a humidified incubator at 37 °C with 10% CO_2_. Transfection of plasmid DNA was performed using Lipofectamine 3000 (Life Technologies) according to the manufacturer instructions.

### Cloning of caspase-2 phospho site mutants and other constructs

All phosphorylation site mutants were generated by QuikChange site-directed mutagenesis of pEGFP-mouse caspase-2 WT^44^ or pcDNA-DEST47-mouse caspase-2-WT (for T161A) as PCR template. All primer sequences are outlined in Supp. Table 1. To produce BiFC-caspase-2-S384E constructs, the caspase-2-S384E was amplified from the pEGFP-capase-2-S384E using the following PCR primers: VN-caspase-2-S384E (forward: 5’-AAAGATCTCGCGGCGCCGAGCGGGAG-3’ and reverse: 5’-AATCTAGA CGTGGGTGGGTAGCCTGG-3′ with added BglII and XbaI site); VC-caspase-2-S384E (forward: 5’-AAAGATCTCCGCGGCGCCGAGCGGGAG-3’ and reverse: 5’-AA*CTCGAGT*CGTGGGTGGGTAGCCTGG-3′ with added BglII and XhoI site). The amplified fragments were cut with BglII/XbaI or BglII/XhoI and subsequently subcloned in-frame into pBiFC-VN173 and pBiFC-VC155 (gifts from Lisa Bouchier-Hayes, Baylor College of Medicine, Houston, TX, USA), respectively. For bacterial expression, pGEX4T3-Caspase-2-WT^23^ was used as a template for PCR-based mutagenesis to introduce the S384E, S384A or C320G mutations, followed insertion into the BamHI and XhoI sites of pGEX4T3. To obtain the small fragment of caspase-2 containing S384 (Casp2_363-423_ -WT or -S384A), the corresponding fragments (aa 363 – 423) were amplified from the pGEX4T3-Caspase-2-WT or -S384A using the following PCR primers: (forward: 5’-AAAGGATCCGACATGATATGTGGCTATGCTTG-3’ and reverse: 5’-AAACTCGAGTTAGCCAGGGGCATAG CCTTC-3′) and subcloned into pGEX-4T-3 using BamHI and XhoI sites. To obtain human AURKB expression plasmid construct, full-length AURKB was amplified from cDNA made from U2OS cells using the following PCR primers: (forward: 5’-AAACATATGGCCCAGAAGGAGAACTCC-3’ and reverse: 5’-AAACTCGAG GGCGACAGATTGAAGGGC-3′) and subcloned into pET-32a vector plasmid via NdeI and XhoI sites. Each construct was confirmed by DNA restriction enzyme digestion and DNA sequencing.

### Expression and purification of recombinant proteins

The plasmids pGEX4T3-caspase-2-WT, -S384E or -S384A, pGEX4T3-Casp2^363-423^- WT or -S384A) or pET32a-AURKB were transformed into *E.coli* BL21(DE3)pLysS (Promega). Single colonies were inoculated into 50 ml LB with 100 µg/ml ampicillin and cultured overnight at 37 °C with shaking at 180-200 rpm. The next day, 10 ml overnight culture was diluted into 400 ml LB with ampicillin (100 µg/ml) and cultured at 30 °C. To induce recombinant proteins isopropyl-β-D-thiogalactoside (IPTG, Sigma-Aldrich) was added (0.1 mM) when OD_600_ was between 0.4 – 0.6 and the culture was incubated for 24 hours at 16 °C. Bacterial cells were harvested by centrifugation at 5,000 rpm for 10 mins at 4 °C. For GST-tagged recombinant protein purification, the pellet was resuspended in 20 ml of cold lysis buffer (25 mM HEPES, 10% sucrose, 0.1% CHAPS, 1 mM EDTA, 10mM DTT, pH 7.4) with protease inhibitor (Sigma-Aldrich), followed by sonication (10 x 20- second pulses). Triton X-100 (Sigma-Aldrich) was then added to a final concentration 0.1 % and incubated for 1 hour at 4 °C with rocking, followed by centrifugation at 15,000 rpm for 30min at 4 °C. Supernatant was transferred to a new 50 ml falcon tube and 250 µl bed volume of PBS-washed glutathione Sepharose beads (GE Healthcare Australia) was added to the lysate, followed by overnight at 4 °C with rocking. Supernatant containing beads were transferred to a column in a 4 °C cold room and flow-through was collected. After washing beads four times with 10 ml of PBST (PBS, 0.1% Triton X-100). GST-tagged recombinant protein was eluted in 10 bead volumes of elution buffer (50 mM Tris, 10 mM reduced glutathione, pH 8.0) in 10 elute fractions. For AURKB purification, bacterial pellet was resuspended in 20 ml of cold His-lysis buffer (50 mM NaH_2_PO_4_, 300 mM NaCl, 10 mM imidazole, pH 8.0) with protease inhibitor (Sigma-Aldrich), followed by sonication (10 x 20-second pulses), followed by centrifugation at 15,000 rpm for 30min at 4 °C. The supernatant was transferred to a new 50 ml falcon tube and 250 µl bed volume of Ni-Sepharose (GE Healthcare Australia) was added to the lysate, followed by overnight at 4 °C with rocking and transferring to a column in the cold room. After washing beads four times with 10 ml of wash buffer (50 mM NaH_2_PO_4_, 300 mM NaCl, 20 mM imidazole, pH 8.0), His-AURKB was eluted in 10 bead volumes of elution buffer (50 mM NaH_2_PO_4_, 300 mM NaCl, 250 mM imidazole, pH 8.0) in 10 elute fractions. Purified proteins were checked by SDS-PAGE analysis. Pooled eluates were dialysed against 50 mM Tris-HCl, pH 7.5. Protein concentration was quantitated with BCA Protein Assay (Thermofisher Scientific) and stored in aliquots at −80 °C until used.

### Protein pull-down assay

To examine the direct interaction between caspase-2 and AURKB, GST pull-down assay was carried out. GST or GST-Casp2-C320G (1 µg) was incubated with 500 ng His-AURKB in 300 µl incubation buffer (50mM Tris-HCl, 10mM MgCl2, 0.1 mM EDTA, 0.01% Brij 35, pH7.5) for 2 hours with rotation, at room temperature. This was followed by addition of 10 µl glutathione-agarose beads (GE Healthcare) and further incubation for 1 hr with rotation at room temperature. After four washes with incubation buffer, beads were heated in 2 x Laemmli buffer for 5min at 95 °C. Samples and 10% inputs were then subjected to SDS-PAGE and immunoblotting.

### Purification of caspase-2-C320G-GFP

U2OS-*CASP2^-/-^* cells were seeded into 16 x 100 mm dishes at a total density of 1.5 x 10^6^ cells/dish. Cells were treated with 20 µM Z-VAD for 2 h and then transfected with 6 µg pEGFP-caspase-2-C320G for 48 h. The cells were harvested and lysed in RIPA buffer [25 mM Tris-HCl pH 7.4, 150 mM NaCl, 1% nonyl-phenoxylpolyethoxylethanol (NP-40), 1% sodium deoxycholate, 0.1% sodium dodecyl sulphate (SDS)] supplemented with 1X Halt Protease and Phosphatase Inhibitor Cocktail, EDTA (Thermo Scientific), and 10 µM N-Ethylmaleimide (Sigma-Aldrich). Lysed cells were sonicated and lysates cleared by centrifugation at 13,200 rpm (4 °C). GFP-Trap (ChromoTek) was used to purify caspase-2-C320G-GFP, according to the manufacturer’s instruction. Briefly, 150 µl GFP-Trap_MA bead suspension was equilibrated by washing with RIPA buffer three times. Cell lysates were added to the GFP-Trap_MA beads and incubated at 4 °C overnight with gentle rocking. The beads were then separated magnetically until supernatant was clear. The supernatant was collected as ‘flow-through’ sample for SDS-PAGE and Western blot analysis. The beads were resuspended in 1 mL ice-cold wash buffer (10 mM Tris-HCl, 150 mM NaCl, 0.5 mM EDTA, pH 7.5), briefly spun and magnetically separated. The supernatant was removed, and washing was repeated four more times with high salt wash buffer (10 mM Tris-HCl, 500 mM NaCl, 0.5 mM EDTA, pH 7.5). Bound GFP-caspase-2-C320G was eluted by adding 50 µl elution buffer (0.2 M glycine pH 2.5), followed by incubation for 2 min and magnetic separation. The supernatant was transferred to a new tube and 5 µl of 1M Tris (pH 10.4) was added for neutralisation. Elution was repeated three times. A sample of each fraction (10 µL) was subjected to SDS-PAGE analysis and Coomassie blue staining (Supp. Fig. 1). ‘Elution 1’ was utilised for proteomic analysis.

### Mass spectrometry and data analysis

Protein samples were resuspended in 6M Urea, 100 mM DTT and 100 mM Tris-HCl pH7.0 and subjected to protein digestion using FASP (filter aided sample preparation)^45^. Peptides were collected and acidified with formic acid (FA) to a 1% final concentration. Solvent was removed in a CentriVap concentrator (Labconco) and peptides were resuspended in MilliQ water containing 1% acetonitrile (ACN) and 1% formic acid. Samples were analyzed by nanoflow liquid chromatography tandem-mass spectrometry (LC-MS/MS) on a nanoAcquity system (Waters, Milford, MA, USA) coupled to a Q-Exactive mass spectrometer (Thermo Fisher Scientific, Bremen, Germany) through a nanoelectrospray ion source (Thermo Fisher Scientific). Peptide mixtures were directly loaded onto a 250 mm column with 75µm inner diameter (nanoAcquity UPLC 1.7µm BEH130 C18) on a 120 minute linear gradient from 1% to 35% buffer B (A: 99.9% Milli-Q water, 0.1% FA; B: 99.9% ACN, 0.1% FA) at a 400nl/min constant flow rate. The Q-Exactive was operated in a data-dependent mode, switching automatically between one full-scan and subsequent MS/MS scans of the ten most abundant peaks. The instrument was controlled using Exactive series version 2.6 and Xcalibur 3.0. Full-scans (m/z 350–1,850) were acquired with a resolution of 70,000 at 200 m/z. The 10 most intense ions were sequentially isolated with a target value of 10000 ions and an isolation width of 3 m/z and fragmented using HCD with normalized collision energy of 27 and stepped collision energy of 15%. Maximum ion accumulation times were set to 50ms for full MS scan and 200ms for MS/MS. Underfill ratio was set to 2% and dynamic exclusion was enabled and set to 60 seconds.

The raw files were analysed using the MaxQuant^46, 47^ software (version 1.5.8.3), The database search was performed using mouse protein sequences obtained from Uniprot including isoforms with strict trypsin specificity allowing up to 2 missed cleavages. The minimum required peptide length was set to 7 amino acids. Carbamidomethylation of cysteine was set as a fixed modification while N-acetylation of proteins N-termini, oxidation of methionine, phosphorylation of S/T/Y and GlyGly on lysine were set as variable modifications. During the MaxQuant main search, precursor ion mass error tolerance was set to 4.5 ppm and fragment ions were allowed a mass deviation of 20 ppm. PSM and protein identifications were filtered using a target-decoy approach at a false discovery rate (FDR) of 1%.

### Caspase-2 dimerisation assay

Bimolecular fluorescence complementation (BiFC) analysis for caspase-2 activation was performed as described previously with minor modification^48^. U2OS-*CASP2^-/-^* cells were seeded onto 13 mm glass coverslips (Thermo Fisher Scientific) and incubated overnight at 37 °C in a 10 % CO_2_ incubator. The next day, the medium was replaced with complete culture medium containing 20 µM Z-VAD-FMK (Sigma Aldrich) 2 h prior to transfection. The cells were then co-transfected with 150 ng of the pBiFC-HA-Casp2(S384E)**-**VC155 and pBiFC-HA-Casp2(S384E)**-**VN173 (mouse caspase-2) for BiFC and 10 ng of pDsRed-Mito (Clontech) as a transfection reporter plasmid, using Fugene HD reagent (Promega). pBiFC-HA-CASP2 (C320A)-VC155 and pBiFC-HA-CASP2 (C320A)-VN173 (human caspase-2) were used as system control ^12^. After incubation for 2 h, DMSO (control), or 100nM BI 2536 (Sigma Aldrich) were added to the medium and cells were incubated a further 24 h at 37°C in a 10 % CO_2_ incubator. Cells were then fixed with 4% paraformaldehyde (Sigma Aldrich) in PBS and BiFC imaged by confocal microscopy using a Leica TCS SP8 (Leica, Wetzlar, Germany). At least 100 cells were counted in five different areas in two independent experiments to quantify BiFC-positive cells.

### Caspase activity assay

Caspase activity assay using VDVAD-AFC was carried out as previously described^49^. The protein concentration of cell lysates were quantified by BCA Protein Assay (Thermofisher Scientific) according to manufacturer instructions and 50µg of cell lysate was mixed with Caspase-2 Activity Buffer (0.1 M MES [2-(N-morpholino) ethanesulfonic acid], 10% sucrose, 0.1% CHAPS, 0.5 mM EDTA, pH 6.5) to a total volume of 50 µL. VDVAD-AFC (100µM, Sigma Aldrich)) diluted in 50 µL Caspase-2 Activity buffer was added to each sample to give a final volume of 100 µL per well. Fluorescence was measured in 10 min intervals for 170 mins at 37 °C, with excitation = 400 nm, emission = 505 nm, to quantitate caspase-2 activity. For GST recombinant proteins, 20 µL of crude *E. coli* extract was used.

### Generation of caspase-2 knockout cell lines

Caspase-2 knockout cell line was generated using CRISPR/Cas9 technology as described previously ^50^. pSpCas9(BB)-2A-Puro (PX459) was a gift from Feng Zhang (Addgene plasmid # 48139 ; http://n2t.net/addgene:48139 ; RRID:Addgene_48139)^50^ The sgRNA oligos (sense: 5’ - caccgaccaaaaatgttcttcatcc - 3’; anti-sense: 5’-aaacggatgaagaacatttttggtc-3’) were annealed, phosphorylated by T4 PNK (NEB, Ipswich, MA, USA) and inserted into BbsI (NEB, Ipswich, MA, USA) site of pSpCas9(BB)-2A-Puro (PX459) vector plasmid. To establish the knockout cell line, 2 x 10^5^ U2OS cells were transfected with 2 µg of sgRNA and Cas9 expressing plasmid using Fugene HD reagent (Promega, Madison, WI, USA) according to the manufacturer’s instruction. pSpCas9(BB)-2A-GFP and pSpCas9(BB)-2A-Puro (PX459) were also transfected as a negative control and positive control, respectively, for puromycin treatment. The cells were treated with 1 µg/ml puromycin for 2 days until all the negative control cells died. The surviving cells were harvested and re-seeded into three 10 cm dishes at 1 x 10^3^ cells/dish for isolating single colonies. To screen candidate single clones, the sgRNA-targeted area was amplified by PCR (forward: 5’-tggtggaagccaactgttgaaacc-3’; reverse: 5’-tctcagaaaggaaggcaaagacacg-3’) and the PCR products were analysed by heteroduplex analysis in PAGE gel as described previously^51^ and DNA sequencing. DNA sequencing results were used for insertion and deletion analysis in the targeted gene area by TIDE software (http://tide.nki.nl). Loss of Caspase-2 expression in the candidate clones was validated by Western blot.

### Generation of stable cell lines immunofluorescence microscopy

U2OS cells stably expressing GFP (sWT) and U2OS-*CASP2^-/-^* cells stably expressing GFP (sKO), GFP-Casp2-C320G (sC320G) or GFP-Casp2-S384E (sS384E) were generated using pEGFP mock, pEGFP-Casp2-C320G or pEGFP-Casp2-S384E plasmids, respectively. For immunofluorescence microscopy, cells were grown on coverslips overnight and fixed in 4% paraformaldehyde in PBS for 10 min at room temperature. The fixed cells were washed in TBS, permeabilized and blocked with blocking solution (3% BSA, 0.1% Triton X-100 in TBS) for 1 h at RT. Samples were incubated overnight with goat anti-GFP antibody at 1:500 dilution in blocking solution. After three washes in TBS/0.1% Triton X-100, coverslips were incubated for 60 min with anti-goat IgG-Alexa568 (Molecular Probes) diluted in blocking solution. Samples were washed another three times (as above) and coverslips were mounted onto glass slides with mounting medium containing DAPI (Life Technologies). Images were taken with a Zeiss LSM-800 confocal microscope (Zeiss).

### Immunoblotting

Protein samples were denatured for 5 min at 95 °C and run on Mini or Midi TGX-Stain-free precast gels (Bio-rad) at 110V for 60 min (mini gels) or at 140V for 60 min (midi gels) in SDS-PAGE running buffer (250 mM Tris, 192 mM Glycine, 0.06% SDS). The proteins were transferred onto PVDF membrane (Bio-Rad) using a Trans-Blot Turbo (Bio-Rad), according to the manufacturer’s instructions, in transfer buffer (25 mM Tris, 192 mM Glycine, 20% methanol, 0.05% SDS). Membranes were then incubated in blocking buffer (TBS-T (20 mM Tris, 150 mM NaCl pH 7.4, 0.05% Tween-20), 5% (w/v) skim milk powder (Diploma)) for 1h at RT, followed by incubation with primary antibody overnight at 4 °C. Following membrane washing in TBST for 4 x 15 min, membranes were incubated with horse radish peroxidase (HRP)-conjugated secondary antibodies for 1hr at RT. Membranes were then washed in TBST for 4 x 10 min, followed by development with Enhanced Chemi-Luminescence (ECL) reagent (Pierce Chemical Co., Rockford, IL). Fuji LAS4000 System (GE Healthcare) or Chemidoc MP (Bio-Rad) were used for detecting luminescence signals.

### DNA content analysis

DNA content analysis was performed as described previously ^14^. Briefly, harvested cells were washed with ice-cold 1 X PBS followed by fixation in 70% ice-cold ethanol in PBS. Fixed cells were incubated overnight at −20 °C. Cells were centrifuged at 1200 rpm for 5 mins then resuspended in 2 mL 1X PBS for rehydration. Cells were centrifuged a second time and pellets resuspended in 1 mL 0.25 % Triton X-100 (Sigma Aldrich) in PBS. Following a final centrifugation step, cell pellets were resuspended in 400 µL staining solution [25 µg/ml propidium iodide (Sigma Aldrich), 40 µg/ml RNase A (Sigma Aldrich)] and incubated for 2 h at room temperature in the dark. The stained cells were stored at 4 °C until flow cytometric analysis on a Fortessa (BD Biosciences).

### Cell death and viability assay

For cell death assays, WT or *Casp2^-/-^* immortalised MEFs ^11^ were co-transfected with GFP-caspase-2-WT, - S384A or -S384E and β-gal expressing plasmids. After 24 h, cells were fixed and incubated with an X-gal containing solution. Blue (transfected) cells were counted for apoptotic morphology with a stereomicroscope (Nikon, Tokyo, Japan). At least 400 cells were counted in at least 20 different fields for each construct. Cell viability was determined using the MTS assay. Briefly, cells were seeded in triplicate at 5000 cells per well in 50 µL complete culture media in a 96-well microplate (Falcon) and cultured overnight at 37 °C with 10% CO_2_. Culture medium (50 µl) containing DMSO or 200 nM BI 2536 was added, followed by 48 h incubation at 37 °C with 10% CO_2_. Twenty-five microliters of MTS/PMS (96:4) reagent was added to each well. The plate was then incubated in a humidified, 10% CO_2_ incubator for 4 h, followed by reading absorbance at 490 nm using a FLUOstar Omega (BMG Labtech, USA). Control wells (no cells), were used to detect the cell-free background absorbance.

### *In vitro* kinase assay

*In vitro* kinase assays were performed as described previously^52^. Proteins were incubated with 38 ng of Aurora B kinase (PV6130 LifeTechnologies) or 125 ng of Aurora A kinase (PV3612 LifeTechnologies), 0.1mM ATP (Sigma-Aldrich), 5 microcuries (μCi) of [γ-^32^P]-ATP (3000 Ci/mmol, 10 mCi/ml) (PerkinElmer) in kinase buffer (20mM HEPES pH 7.5, 2mM MgCl_2_, 1mM DTT) in a final reaction volume of 12.5 μl. After 30 min incubation at 30°C with constant agitation, 12.5 μl of 2x Laemmli sample buffer were added to stop the reaction. Samples were heated for 10 min at 90°C and loaded on a 4–20% Tris-Glycine precast gel (ThermoFisher). Protein was transferred onto a nitrocellulose membrane using the iBlot Dry Blotting System. Membranes were then exposed to Kodak BioMax XAR Films (Sigma-Aldrich) at −80°C. The membranes were then blocked and subjected to immune blotting (IB) for GST.

### Statistical Analysis

The data are presented as mean ± standard error of the mean (SEM) and were considered statistically significant when p < 0.05. A two-sided Student’s t-test was used to analyse differences between data groups unless otherwise stated. All statistical analyses were conducted using GraphPad Prism, Version 6.05 (GraphPad, supplied by University of South Australia; Graph Pad Inc., CA).

### Data Availability

The mass spectrometry proteomics data have been deposited to the ProteomeXchange Consortium via the PRIDE [1] partner repository with the dataset identifier PXD017866, and are available via ProteomeXchange.

## Acknowledgments

We thank J Puccini for helping with the U2OS-*CASP2^-/-^* cell line, CCB Cytometry for DNA content analysis, and members of our laboratory for discussions and useful comments. This project was supported by the National Health and Medical Research Council (NHMRC) of Australia project grants 1043057 and 1156601, a NHMRC Senior Principal Research Fellowship (1103006) and a University of South Australia support package to SK. We acknowledge the support of the Australian Cancer Research Foundation funded Cancer Genomics and Imaging Core Facilities at CCB. JMM acknowledges the NHMRC fellowship (1105754, 1172929) support; JJS, AIW and JMM acknowledge NHMRC IRIISS (9000587); and the work in PPD laboratory is supported by a BBSRC grant (BB/R001227/1).

## Author Contributions

YL designed and performed experiments, analysed data and wrote the paper; DDB helped with the generation of mutants and performed cell biology experiments; JS and AW carried out MS analysis; LC and PD performed *in vitro* phosphorylation and immunoblotting experiments; JM generated the caspase-2 structure modelling; LD and SK designed and supervised the study, analyzed data, acquired funding and wrote the paper. All authors discussed the results and commented on the manuscript.

